# Distribution of Critical Areas for Ecological Conservation in Multiple Scenarios——Emphasizing the Impact of Human Activities

**DOI:** 10.1101/2022.03.07.483357

**Authors:** Xinyan Dai, Hongwei Wang, Chen Ma, Xiaoqin Wang, Jing Zhou, Bo Tan

**Author notes:** (X.D.); (C.M.); (X.W.); (J.Z.); (B.T.).

## Abstract

Determining critical ecological protected areas at the city (county) spatial scale is crucial for coordinating regional ecological environment management, control, and governance. It is a recognized consensus in academia that human activities significantly impact natural ecosystems. Many scholars ignore this point in the research process and only use several factors to characterize human influence. Therefore, this study takes Aksu City and Wensu County, important cities and towns in southern Xinjiang, as examples, focusing on the impact of human activities on the distribution of critical ecological protection areas. First, we simulated the range, intensity, and local natural conditions of human activities in the study area using geospatial data. We created corresponding resistance surfaces (human activity resistance surface and natural landscape resistance surface). We then assign different weights to the two resistance surfaces based on several possible scenarios, resulting in different synthetic resistance surfaces. Finally, we used the Linkage Mapper plugin to identify critical ecological reserves and compared several results. The results show that: Human activities have greatly interfered with the migration and dispersal of species, affecting the distribution of critical ecological reserves. The specific performance is that in the urban center area with high intensity of human activities, the number and location of the surrounding ecological corridors change significantly with the increase in the proportion of human activity resistance and the total area remains unchanged; As part of the ecological corridors, the ecological pinch points changes with the location of the corridor, and the whole area remains unchanged; The threshold range of the restoration value of ecological barrier points is reduced. The maximum value generated by the restoration of ecological barrier points is reduced, which shows that the restoration value of ecological barrier points decreases with increased human activities.

## 1 INTRODUCTION

Countries worldwide gradually realize that the destruction of the ecological environment will bring severe consequences and begin to place ecological protection in an important strategic position. Ecological security has received more and more attention at home and abroad and has become an essential scientific issue that all countries must face together and urgently need to solve (Schreurs M A,1998). With the rapid and irrational development of human society in the past few hundred years, especially in the mid-to-late 50 years of the 1990s, the disturbance of human activities to the natural ecosystem has been increasing, and the threat to the ecological environment has been increasing. Human activities have become the dominant-negative factor affecting the stability of the natural environment in various circles on the earth (World Resources Institute, 2005). Some studies also show that human activities profoundly affect the behavior of wild animals (Nadal J,2022; Rutz C,2020; Stillman J H,2019; Riotte-Lambert L,2020; Nickel B A,2020). The establishment of ecological corridors is an important measure to solve the ecosystem’s fragmentation caused by the current violent human activities and the many ecological and environmental problems that follow (Rosenberg D K,1997; Gregory A,2021). Ecological corridors are priority areas for ecological protection (Huang L,2021) and are also an essential component of the ecological security pattern. Establishing an ecological security pattern is a new idea to solve regional ecological security problems effectively (Wang C,2020).

In the early 1990s, the research on this issue in western countries mainly revolved around the concept of ecological security, the interrelationship between ecological security and national security (Hermann C F,1909), sustainable development (Kothari A,1915), and globalization (Amirnejad H,1922). Subsequently, some European countries conducted a lot of research and discussion on the relationship between environmental change and security. In the late 1990s, some results were obtained, such as “Environment and Security in the International Context” in 1999 (Lietzmann K M,1999). In the 21st century, the research gradually penetrated the internal factors affecting environmental safety, which promoted the development of related theories and indicators (Zhen-li L U O.2002). For the study of ecological security patterns, the west started earlier. As early as the end of the 19th century, Frederick Law Olmsted used natural elements such as rivers as a link, using open native greenery to connect several parks into a landscape belt that stretched about 16 kilometers. - Emerald necklace. The landscape belt has long been regarded as a successful example of the protection and application of the regional ecological security pattern (Runte, A,1998). In 1984, the concept of Ecological Infrastructure (EI) in the study of UNESCO’s “Man and the Biosphere Program” (MAB) aimed to promote the coordinated development of ecologically sustainable development and cities (UNESCO,1984). This concept was subsequently applied in biological conservation to optimize the design of habitat networks (GG Bruinderink,2003), emphasizing its role in providing biological habitats (Ahern J. 1995) and producing energy resources.

Since the 18th National Congress of the Communist Party of China, vigorously promoting the construction of ecological civilization has been incorporated into China’s national strategic goals. The report of the 19th National Congress of the Communist Party of China in 2017 pointed out that it is necessary to “intensify the protection of ecosystems, optimize the ecological security barrier system,” and establish and improve the national land space development and protection system. In May 2020, the Ministry of Natural Resources issued the “Guidelines for the Compilation of Urban Territorial Space Overall Planning (Trial).” This guide clearly proposes building a continuous, complete, and systematic pattern of ecological protection and prioritizing the delineation of protection space. Ecological security models are essential for maintaining or controlling some ecological processes in an area. It plays a fundamental role in ensuring the security of regional ecological space. Therefore, identifying the critical areas of ecological protection and building a scientific and reasonable ecological protection pattern is a severe challenge facing the current ecological space protection and restoration project. It has also become a hot direction for domestic scholars. Research on ecological security patterns in China involves habitat protection (Bai L,2019), urban planning (Su Y,2016), biodiversity conservation (Zhao X Q,2015), and other fields, including urban agglomerations (Ztl A,2020; Guo R,2019; Wang D, 2019), provincial areas (Li J, 2019; Sun M, 2020), counties (Zhao X, 2021; Fu Y, 2020) and other scales. Methods such as ecosystem service evaluation (Peng J, 2018; Zou C, 2018) or habitat quality evaluation (Wang Y, 2019) are usually used in the selection of ecological sources. When constructing ecological corridors between important ecological landscapes, methods such as the least-cost path (Lin Q,2016), circuit model (Gao J,2020), and other methods are often used together with the landscape spatial morphology method (An Y, 2020). Some scholars have also improved the model (Li S, 2019) to meet the research needs. It is worth mentioning that some scholars only consider the landscape resistance when constructing the ecological resistance surface (Zhao S,2019) or simplify the human activity resistance to one or several factors (Li H,2010). This approach hardly reflects the impact of human activities on wildlife migration. Therefore, when constructing the ecological resistance surface, the influence of the natural landscape and human activities on species expansion and migration must be comprehensively considered. We use enterprise nuclear density analysis data, road line density analysis data, annual average luminous brightness value data, and PM2.5 annual average concentration data to construct the human activity resistance surface, which is weighted and superimposed with the natural landscape resistance surface. In addition, considering the unknownness of the specific situation, for the convenience of comparison, this study proposes four possible scenarios and conducts a comparative analysis of the obtained results, aiming to explore the impact of different degrees of human activities on key areas of ecological protection.

## 2 STUDY AREA AND DATA SOURCE

### 2.1 Overview of the study area

The study area belongs to the Aksu area of Xinjiang Uygur Autonomous Region. The Aksu area is between 30°30′ - 42°15′ north latitude and 79°28′ - 81°30′ east longitude, with a total area of about 29,019.3 square kilometers and an east-west span of about 199 km, about 219 km from north to south. The study area is deep in the Eurasian continent on the northwestern edge of the Taklimakan Desert, far from the ocean, and has a typical warm temperate continental arid climate. The climate is suitable, and the terrain is flat, the land is fertile, the light and heat are sufficient, the frost-free period is extended, suitable for the growth of various crops. Its territory is rich in wildlife resources. However, with the increase in human population and large-scale land reclamation over the past few decades, the habitat has been dramatically reduced. The number of wild animals has been dramatically reduced. Since the 18th National Congress of the Communist Party of China, the Aksu area has been carrying out ecological civilization construction for many years, and various measures to improve the ecological environment have achieved initial results. Since the 18th National Congress of the Communist Party of China, the Aksu region has carried out ecological civilization construction for many years. Various measures to improve the ecological environment have achieved initial results. Wild camels, egrets, snow leopards, and other precious wild animals have reappeared in the public eye (Hong Wen L,2021).

### 2.2 Data and its sources

This article mainly uses three kinds of data: vector, raster, and text.

#### Vector data

The administrative boundary data and land use data of Aksu City and Wensu County are from the Resource and Environmental Science Data Center of the Chinese Academy of Sciences (http://www.resdc.cn), with a resolution of 30m*30m;

#### Raster data

The average annual vegetation net primary productivity (NPP) data in 2020 obtained through the calculation of the CASA light energy utilization model, from the Geographical National Conditions Monitoring Cloud Platform (http://www.dsac.cn), with a resolution of 500m* 500m; The digital elevation data comes from the Geospatial Data Cloud website (http://www.gscloud.cn) with a resolution of 30m*30m;

The average Chinese nighttime light data in 2020 is from EARTH OBSERVATION GROUP (https://payneinstitute.mines.edu/), with a resolution of 500m*500m;

The spatial distribution data of the average annual PM2.5 concentration in 2019 simulated by kriging interpolation, with a resolution of 1km*1km;

#### Text data

The enterprise data with exact addresses comes from the official website of Tianyancha (https://www.tianyancha.com), and the data acquisition time is November 2021.

## 3 RESEARCH METHODS

### 3.1 Habitat quality assessment

This study plans to use the Habitat Quality model in the InVEST model, which evaluates the habitat distribution under different landscape patterns according to the relationship between land use cover types and threat sources, and the response of different habitats to threat sources. It can also reflect ecosystem diversity and its potential level to provide living conditions for species. The data required by this module is easy to obtain and can replace complex methods such as species surveys to quickly evaluate changes in habitat quality and species numbers and determine the priority of conservation. Therefore, this method is beneficial in the absence of species distribution data.

### 3.2 Natural breakpoint method

First, we used the five-level natural breakpoint method to divide the habitat quality into five levels. Then, we slightly adjusted one of the critical values according to the distribution of the grid values of the habitat quality evaluation results. We take areas with habitat quality greater than or equal to this threshold as foreground data, and the rest as background data. Finally, we perform MSPA analysis and use the core area as an ecological resource. Its ecological significance lies in large natural patches, wildlife habitats, forest reserves, etc.

### 3.3 Selection and Interpretation of Resistance Factors

#### 3.3.1 Selection of resistance factors

This study selected seven natural landscape resistance factors based on reference to related research, including land use, vegetation net primary productivity (NPP), slope, slope aspect, terrain relief, elevation, and water area. Species’ needs for habitat quality, water, and energy access. We also consider temperature and topographic constraints on species expansion and migration. Then we considered that various human activities might cause some disturbance to species expansion and migration. We use the 2020 annual average nighttime light intensity data, enterprise nuclear density data, road linear density analysis data, and PM2.5 annual average concentration data as the impact factor for human activities.

Among them, the annual average nighttime light intensity data is used to characterize the range and intensity of human living, commuting, leisure, and other activities.The enterprise kernel density data can be used to characterize the range and strength of human economic production activities due to its distance and density attributes. The road line density analysis not only has distance and density attributes but also includes grade attributes, which is used to indicate the degree of road obstruction to the connection of ecological sources. The primary sources of PM2.5 in the study area are the sand and dust weather in spring and the coal-fired source for heating in winter (Xu Dajian,2016). Most of the exhaust gas emissions contain toxic substances such as heavy metals. Compared with water and soil pollution, air pollution is easy to quantify and inevitable spillover. Therefore, it is more reasonable to use air pollution to characterize the scope of pollution caused by various human activities.

#### 3.3.2 Interpretation of resistance factors

1. Land cover data: The score is determined according to the land cover type and habitat suitability.
2. Net primary productivity of vegetation (NPP): The net primary productivity of natural vegetation results from the interaction between the biological characteristics of plants and external environmental factors. It is an important indicator used to evaluate the structural and functional characteristics of ecosystems and the population carrying capacity of the biosphere.
3. Elevation (DEM): It is a discrete mathematical expression of the topography of the earth’s surface. We selected the ASTER GDEM 30m resolution digital elevation data from the Geospatial Data Cloud website. This data has the advantages of free access, comprehensive coverage, and high precision and is widely used in geoscience-related fields.
4. Slope: the degree of steepness of the surface unit generated by using the elevation data based on the ArcGIS 10.2 platform.
5. Slope aspect: The slope aspect is defined as the direction of the projection of the slope regular on the horizontal plane (it can also be understood as the direction from high to low). The slope aspect has a more significant effect on the mountain ecology, and the orientation of the mountain impacts the sunshine hours and solar radiation intensity. Therefore, we only extract the mountain aspect for normalization.
6. Topographic relief: also known as relief, relative relief, or relative height, it is the maximum relative elevation difference per unit area, which can reflect the relative height difference of the ground, and is a quantitative index to describe the landform.
7. Distance from the water area: We use the Euclidean distance to calculate the distance from each grid cell to the nearest water area based on the ArcGIS 10.2 platform. Considering that the water area has a hindering effect on the expansion and migration of species, the area where the Euclidean distance is 0, that is, the water surface resistance value, is set to the maximum.
8. Light intensity at night: The sensors on the satellite can detect the lights, fire, and other information of the earth at night, which to a certain extent reflects the degree and scope of human activities at night so that this data can be used as a good representation of human activities.
9. Enterprise location core density: Economic production activities are one of the most important activities of human beings. The value of each grid cell in the enterprise kernel density analysis results represents the degree of influence of all enterprises in the area on this grid cell.
10. Annual average concentration of PM2.5: The average concentration of PM2.5 refers to the average content of inhalable lung particulate matter per cubic meter of air. The higher the concentration value, the more serious the air pollution.
11. Road line density analysis: Input the road classification data into the line density analysis tool. The value of each grid cell in the obtained results represents that the cell is affected by roads of different grades within a fixed range.

### 3.4 Scenario Analysis

Since there is insufficient data and related research to show the specific degree of human activities’ influence on species expansion and migration, we set up four scenarios according to possible situations. We made corresponding comprehensive resistance surfaces according to different scenarios:

1. The process of animal expansion and migration in the study area will not be influent by human activities;
2. The influence of natural landscape on animal expansion and migration in the study area is more significant than that of human influence;
3. The impact of natural landscape on animal expansion and migration in the study area is equivalent to the impact of human activities on it;
4. The influence of natural landscape on animal expansion and migration in the study area is less than human influence.

### 3.5 Extraction of ecological corridors, ecological pinch points, and barrier points

Linkage Mapper is a GIS tool to support regional wildlife habitat connectivity analysis. The tool can plot core habitat vector patches and resistance rasters to map the least cost linkages between core areas. The cost-weighted analysis accumulates a graph of total movement resistance as animals leave a specific core habitat. These scripts can standardize and merge corridor maps and form a comprehensive corridor map.

This study uses the “Build Network and Map Linkages” module in the tool to identify and screen ecological corridors in the study area, use the “Pinchpoint Mapper” module to identify ecological pinch points, and use the “Barrier Mapper” module to identify ecological barrier points. We input the superimposed resistance surfaces under the four scenarios respectively and obtain the distribution of the critical areas of ecological protection under the four scenarios.

## 4 ANALYSIS OF RESULTS

### 4.1 Habitat quality analysis

We input the required data into the InVest model, and after calculation, we obtain the habitat quality label image file, which stores the graphic information intact. In order to facilitate the description of the habitat quality results, we adopted the natural breakpoint method, referring to the distribution of grid values, and divided the habitat quality into five grades. The number of grids at each level is obtained by reclassification. See Table 1 for details.

**Tab.1.** Statistics of Habitat quality.

| Habitat quality grade | Number of grid | Proportion | Exponential mean |
| --- | --- | --- | --- |
| Excellent | 997397 | 4.39% | 0.403 |
| Good | 4993994 | 22.00% |  |
| Medium | 4762404 | 20.98% |  |
| bad | 6296903 | 27.74% |  |
| Terrible | 5651276 | 24.89% |  |

According to Figure 3 and Table 1, the overall habitat quality of the study area is relatively low, and the average habitat quality value is about 0.403. The total proportion of habitats with terrible and poor habitat quality was as high as 52.63%. The proportion of habitats with moderate habitat quality was 20.98 %. The sum of the habitats with excellent and good habitat quality was 26.39%. The total area of good and excellent habitats accounted for about 34.16% of the original land cover data. It indicates that part of the habitat was affected by the threat source and degraded. High-quality habitats are mainly distributed north of the study area and scattered in the west and southeast of the study area.

**Fig. 1.**
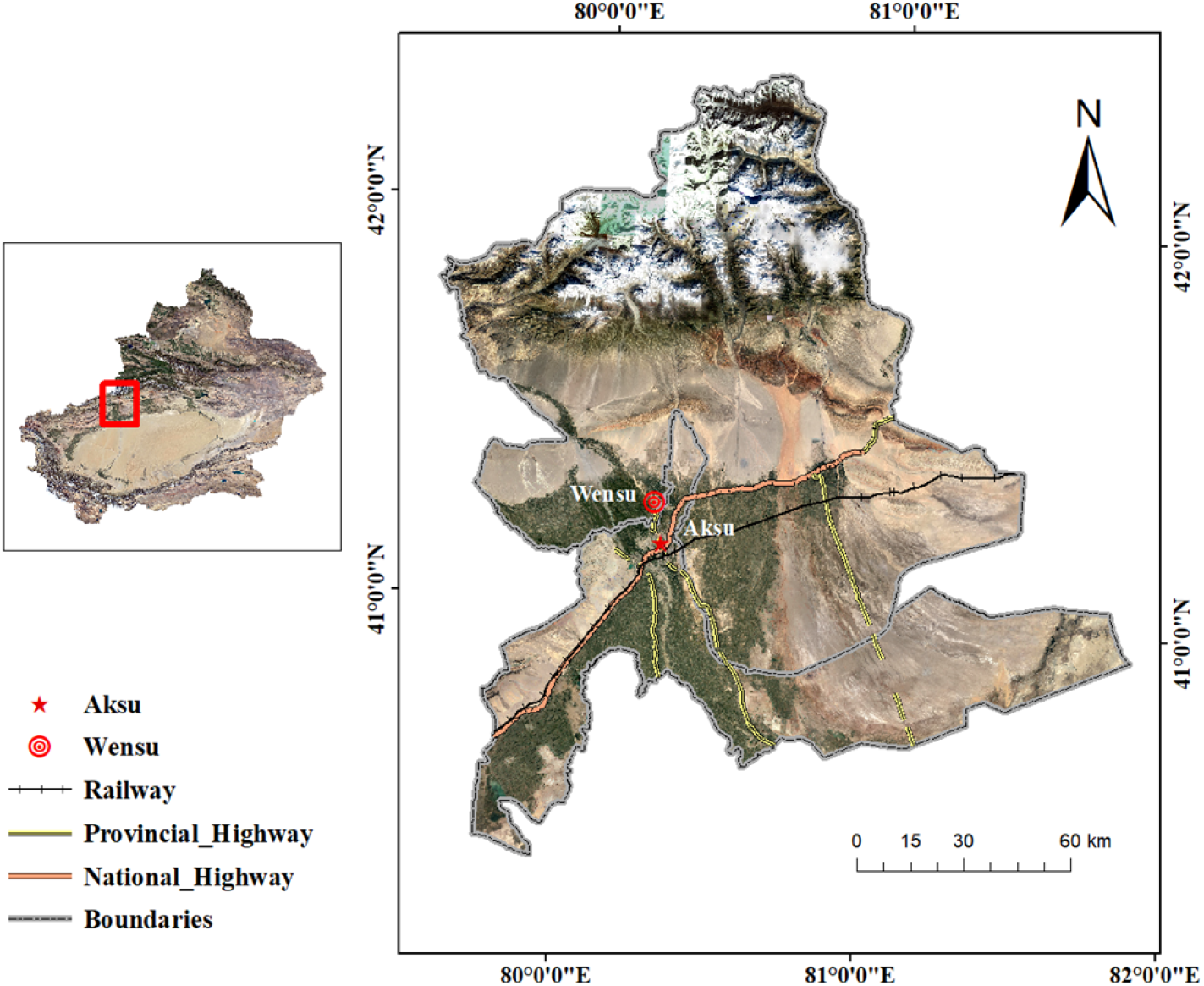
Overview map of the study area.

**Fig. 2.**
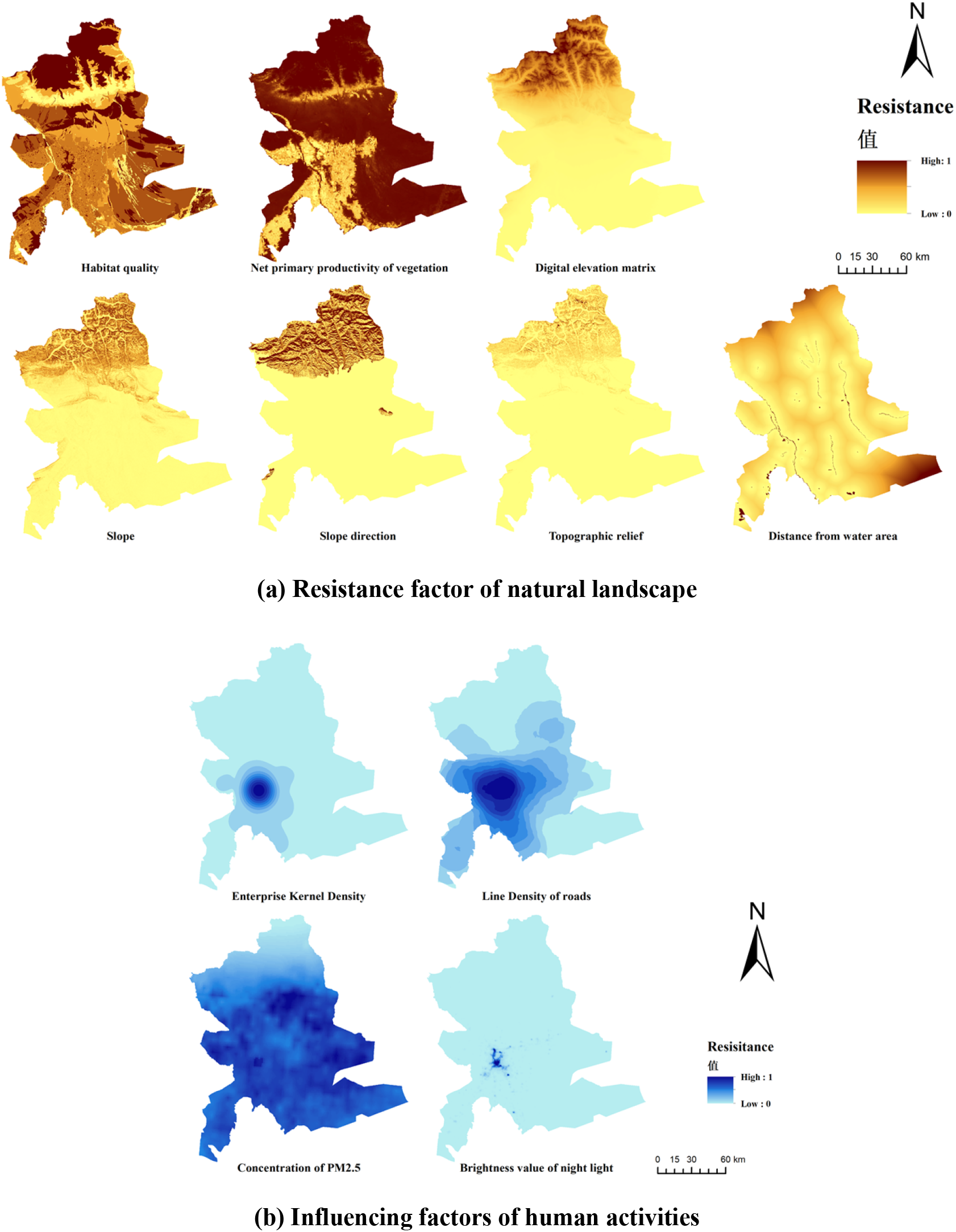
Single resistance factors.

**Fig. 3.**
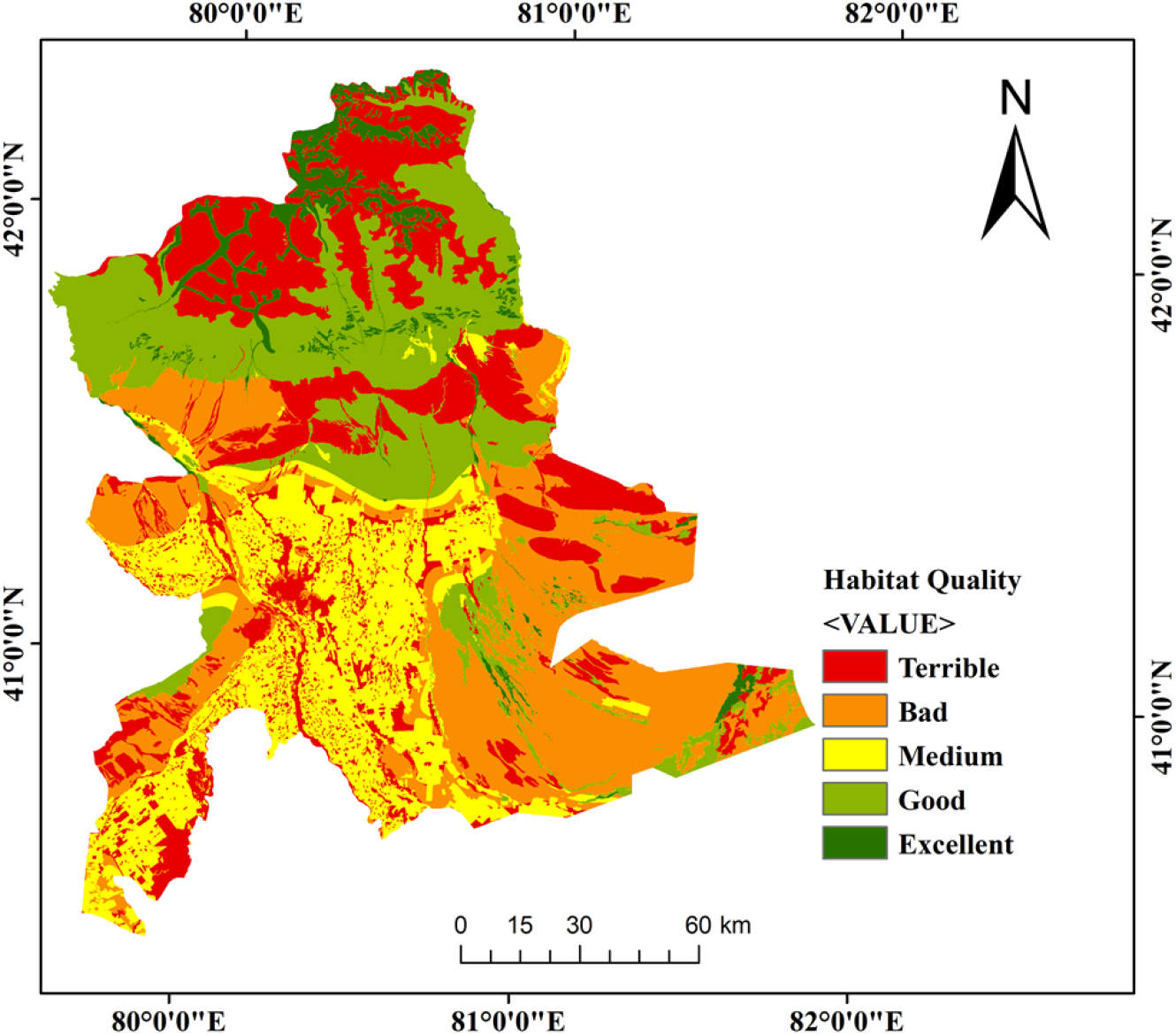
Habitat quality level.

### 4.2 Identification of potential ecological sources

In contrast to the direct use of land-use reclassification results for MSPA analysis, we used a habitat quality assessment model. This model helps us identify and eliminate some of the original land types severely affected by the threat source and cause their habitat quality to decline. This approach improves the accuracy of our identification of ecological sources. (a) Extracted according to land use cover type (b) Extraction according to habitat quality grade

According to Figure 4 and the results of the habitat quality assessment, we found that the foreground patches extracted from the areas with frequent human activities decreased, and the habitat degradation was more pronounced. Ecological sources far from urban centers have not changed significantly. It shows that urban areas where human activities are concentrated have more obvious impacts on habitats.

**Fig. 4.**
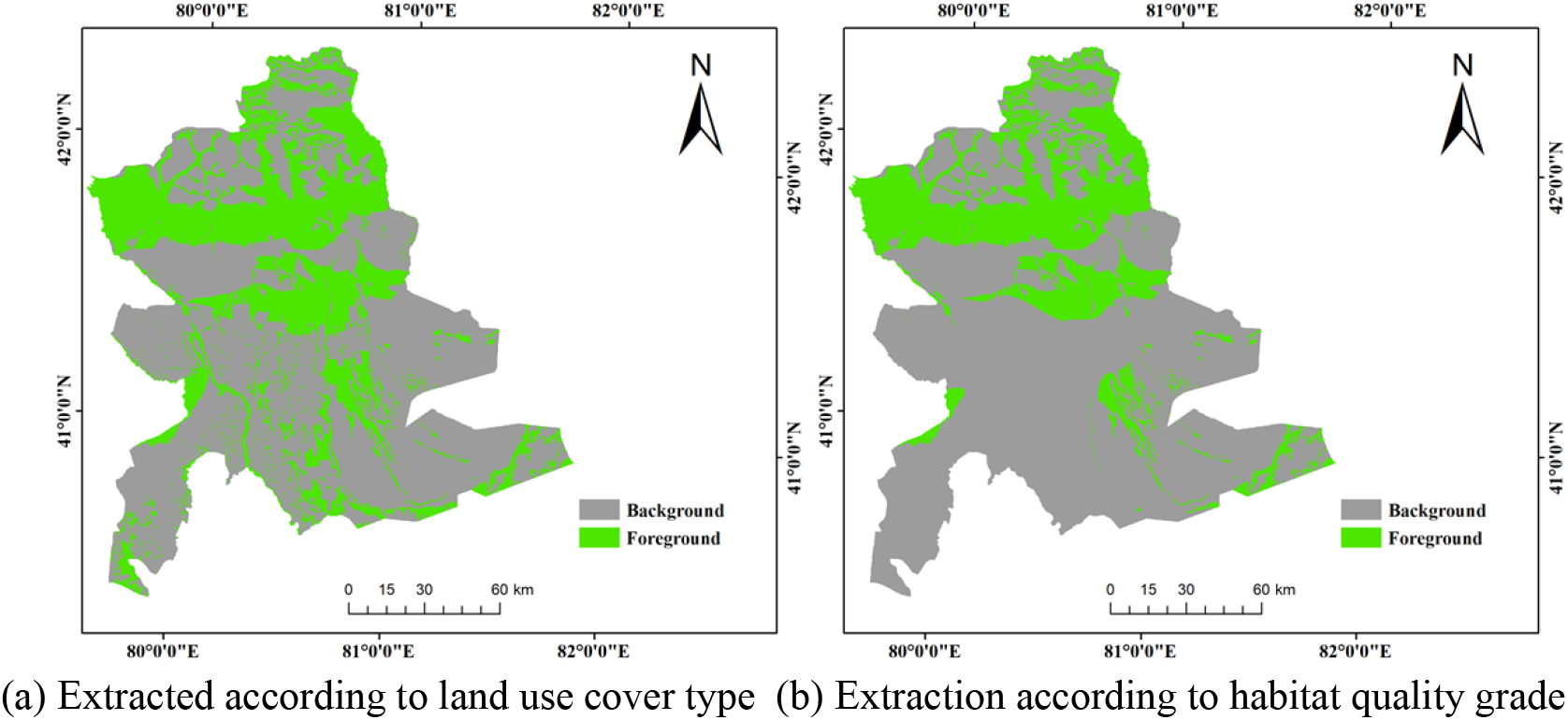
Comparison of foreground data extraction results.

Through MSPA analysis, we extracted 5051.034 km^2^ of the core area. Using the computational geometry function in ArcGIS, ecological patches with more than 5 km^2^ in the core area were selected as ecological sources. The authors selected a total of 28 ecological sources.

### 4.3 Resistance Surface Construction

Each resistance factor is weighted using the Analytic Hierarchy Process (AHP) according to the four possible scenarios mentioned above. All resistance factors are normalized to facilitate overlay processing of multi-source data. Table 3 shows the resistance factors and the weights of each factor under the four scenarios.

**Tab. 2.** Statistics and comparison of Foreground data.

|  | Background grid | Foreground grid | Grid total | Percentage of foreground grid |
| --- | --- | --- | --- | --- |
| Land use | 14948185 | 7753789 | 22701974 | 34.15% |
| Habitat Quality | 16710583 | 5991391 | 22701974 | 26.39% |

**Tab.3.**
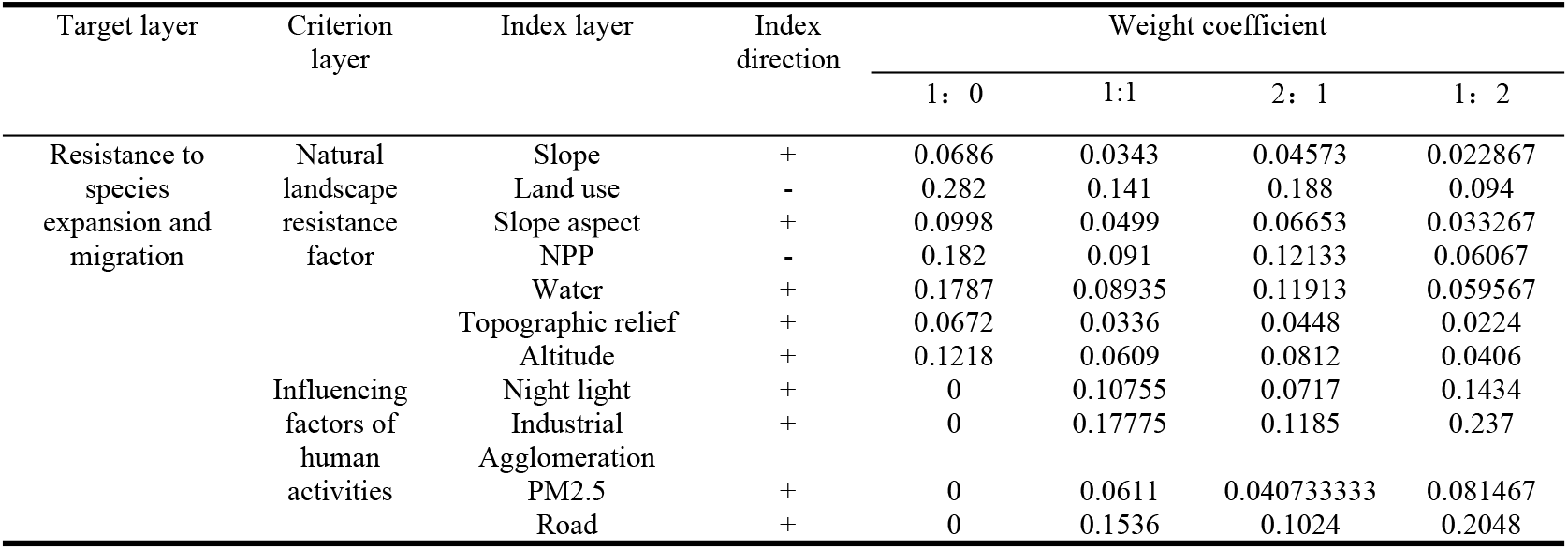
Resistance table.

We considered the four scenarios mentioned above, respectively, according to the sum of the natural local resistance factor weights: the sum of the human interference resistance factor weights is equal to (1) 1:0 (2) 2:1 (3) 1:1 (4) 1:2. Four synthetic resistance surfaces were made using these four scales. In order to meet the conditions of using Linkage Mapper, the author amplifies the resistance surface by 1000 times based on the sum of the weights being 1 to ensure that the measurement methods of the four results are consistent. The results are shown in Figure 5. We can find that as the proportion of human activity resistance increases, the resistance threshold of the comprehensive resistance surface decreases. The high-value resistance area gradually shifted from the east, west, and south periphery to the city center and surrounding areas.

**Fig. 5.**
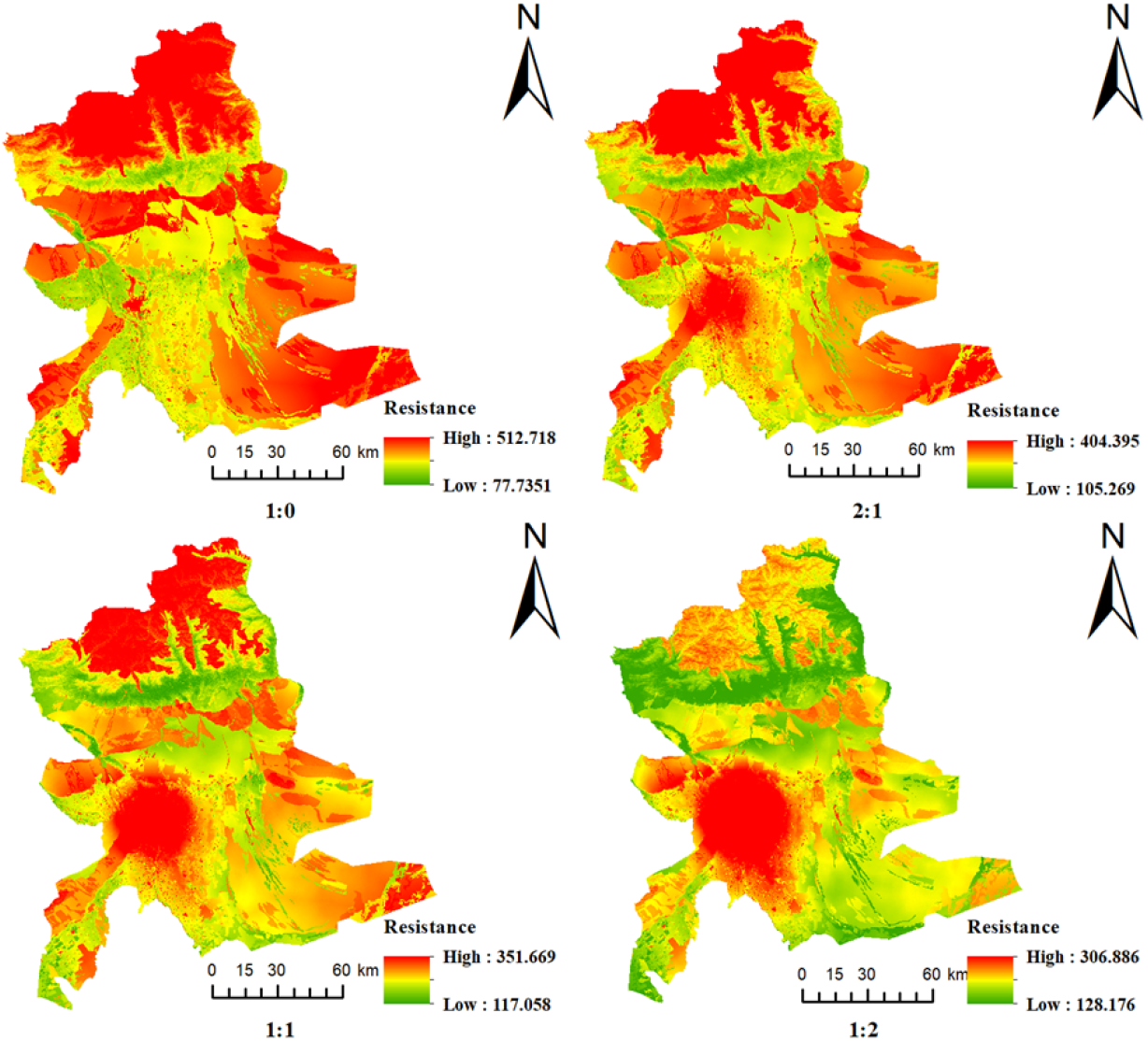
Comprehensive resistance surfaces.

### 4.4 Extraction of ecological corridors, ecological pinch points, and barrier points

The authors input four different resistance surfaces into the Linkage Mapper tool, respectively, to obtain the distribution of critical areas of ecological protection under the four conditions. In order to highlight the ecological pinch points and ecological barrier points and highlight the main changes in the experimental results, the authors zoomed in on the local area and the urban center area.

From Figure 6, we found that two types of ecological corridors, horizontal and vertical, have changed significantly in the area near the town center. The horizontal ecological corridor is located in the middle and south of the town center, and the vertical ecological corridor is located in the northwest direction of the town center:

**Fig. 6.**
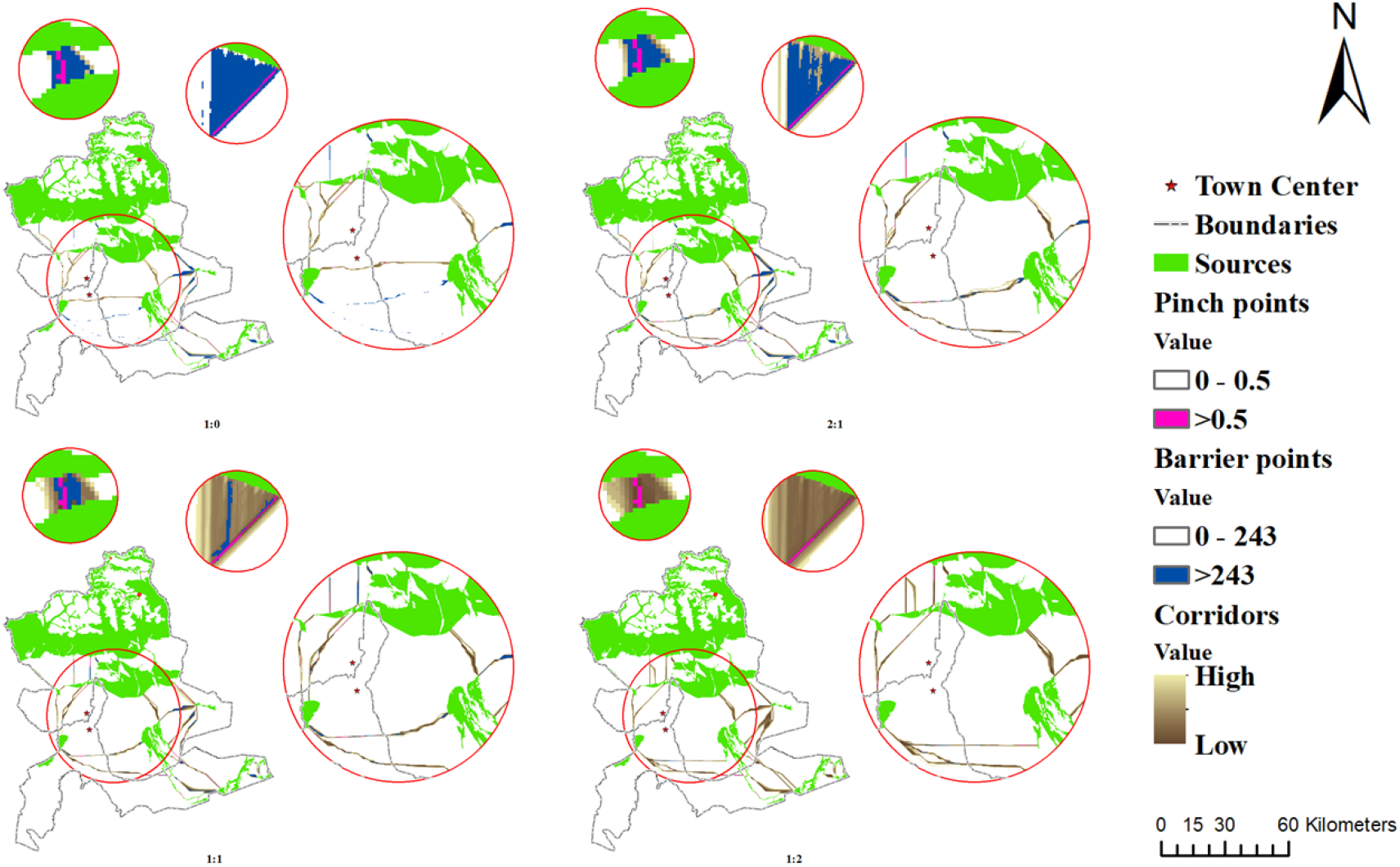
Comparison of key areas of ecological protection.

1. When the ratio of natural landscape resistance and human influence resistance was adjusted from 1:0 to 2:1, the horizontal ecological corridors will increase from the first one to two. A shorter ecological corridor that traverses the town center was replaced by two longer ecological corridors relatively south. It shows that with a slight increase in the impact of human activities, the length of the species migration path increases. At the same time, the branch of the longitudinal corridor decreases by one.
2. When the ratio of natural landscape resistance and human influence resistance was adjusted from 2:1 to 1:1, the horizontal ecological corridors added in (1) expanded outwards. The original vertical ecological corridors also expand outwards. The total length of ecological corridors has increased, but their orientation has not changed significantly.
3. When the ratio of natural landscape resistance and human influence resistance was adjusted from 1:1 to 1:2, the horizontal ecological corridor not only continued to expand outwards, but its trend also changed significantly. The vertical ecological corridor also continued to expand outward, at the same time, a long corridor connecting the north and south ecological patches was lost, and a shorter ecological corridor was added.

In addition, we found that the regional corridors near the town center are long and narrow, and it is more challenging to connect the ecological landscapes. In the areas far from the town center, such as the north and southeast of the study area, the ecological corridors are short and wide, and the ecological landscape is easy to connect.

The ecological pinch point changes as follows:

The ecological pinch points in the study area are presented as narrow strips, smaller dots, and irregular dotted lines. The total area of ecological pinch points is so small that it is difficult to show clearly in Figure 6. With the increase in the proportion of human interference resistance, the area of pinch points with a Pinch point value greater than 0.5 did not change. The location of the ecological pinch near the center of the town moves as the corridor moves.

The changes of ecological barrier points are as follows:

With the increase in the proportion of human activity resistance, the maximum value of repair has experienced a change from 729.196-429.23-373.745-336.917. For the convenience of comparison, we uniformly take the repair value of 243 as the boundary (243 is the median value of the repair value of obstacle points extracted using the purely natural landscape resistance surface). We found that with the increase of the proportion of disturbance resistance by human activities, the area of ecological barriers with restoration value greater than 243 gradually decreased.

## 5 CONCLUSION AND DISCUSSION

### 5.1 Conclusion

This study considers that human activities will interfere with species migration and dispersal, affecting the distribution of critical areas of ecological protection. Therefore, we use several spatial data types to reflect the extent and intensity of human activity. After dimensionless processing, we superimpose different weights of human activity resistance and natural landscape resistance surface to obtain four comprehensive resistance surfaces. Finally, we obtained the distribution of the three critical ecological reserves required to establish the ecological security pattern in the study area under the four scenarios.

By analyzing the above results, we can confirm that the disturbance of human activities has an essential impact on the distribution of critical areas of ecological protection. Especially in the urban center area with high intensity of human activities, the number and location of the surrounding ecological corridors change significantly with the increase of human activity resistance. As a part of the ecological corridor, the location of the ecological pinch varies with the corridor’s location, and the total area remains unchanged. The maximum restoration value threshold of ecological barrier points gradually became smaller, and the total area of ecological barrier points with higher restoration value also gradually decreased. It shows that as the influence of human activities increases, the value generated by the restoration of ecological barrier points decreases.

### 5.2 Discussion

Under the background of the ecological civilization construction strategy, the ecological space planning part of Xinjiang Uygur Autonomous Region’s Territorial Space Planning (2021-2035) has determined the goal of “shaping a green and sustainable ecological space.” At the same time, it also proposed strengthening the protection and control of ecological space, optimizing the system of nature reserves, coordinating the protection and utilization of natural resources, and implementing requirements such as ecological protection and restoration.This study takes Aksu City-Wensu County, an important town in southern Xinjiang, as the research area. We designed a resistance surface that can simultaneously reflect the degree and intensity of human activities affecting the expansion and connectivity of ecological sources. By adding weights to this resistance surface and the natural landscape resistance surface, four comprehensive resistance surfaces are generated, and the critical areas of ecological protection under the four scenarios are determined. When we identified these areas, we considered both the natural context and the disturbance of human activity.

Judging from the evaluation results of habitat quality in the study area, the overall habitat quality of the study area is relatively low. The habitats that can be used as ecological sources are concentrated in the northern part of the study area. The land cover is mainly high-quality forest land, grassland, permanent icebergs, and snow. The ecological sources on the east and west sides and the southeast are scattered, while the urban center area lacks ecological patches that can be used as ecological sources. This situation may be because the northern part of the study area is located in the Tomur Mountains, with high altitude and large terrain fluctuations, and is less affected by human activities. The central and southern parts of the study area are in the plain area, close to the desert, and the vegetation coverage area is small. As the core area of human activities, urban centers are greatly affected by human activities. If we do not intervene, over time, the existing good habitats will inevitably degrade, which will affect the survival of some species.

From the changes of ecological corridors in the experimental results: (1) When the ratio of natural landscape resistance and human influence resistance was adjusted from 1:0 to 2:1, the horizontal ecological corridors increased from the first one to two. Specifically, a shorter ecological corridor that traverses the town center is replaced by two longer ecological corridors relatively south, indicating that the length of species migration paths increases with a slight increase in the impact of human activities. The number of branches in the longitudinal corridor is reduced by one, indicating fewer paths available for species migration. (2) When the ratio of natural landscape resistance to human interference resistance is adjusted from 2:1 to 1:1: both the horizontal ecological corridor and the vertical ecological corridor generated in (1) expand outward, and the total length of the ecological corridor increases, but their direction did not change significantly, indicating that with the further increase of the influence of human activities, the length of species migration path also further increased. (3) When the ratio of natural landscape resistance to human activity resistance is adjusted from 1:1 to 1:2: the horizontal ecological corridor not only continues to expand outwards but also changes its trend significantly. Vertical ecological corridors are also expanding outward, at the same time, the long corridor connecting the north-south ecological patches was lost, and a shorter ecological corridor was added. It shows that when the influence of human activities increases to a certain extent, to avoid being affected by human activities, organisms are bound to abandon the migration paths with obvious natural landscape advantages and open up more difficult migration paths. It undoubtedly dramatically increases the difficulty of species dispersal and is highly detrimental to biodiversity conservation in the study area.

The ecological pinch point is the area with a high current density in the ecological corridor, that is the area where species flow is easiest to pass through and essential to the ecological corridor. In order to facilitate the comparison between the four results, we set the same parameters, so the total area of ecological pinch points does not change; only the location changes with the movement of ecological corridors.

Ecological barrier points are areas with restoration value, that is, the restoration of these areas is of great significance to the connection of ecological plates in the study area. From the change of ecological barrier points: when the ratio of natural landscape resistance to human activity resistance is adjusted from 1:0 to 2:1, many continuous ecological barrier points in the south of the town center are transformed into two important connecting corridors. These two ecological corridors can connect ecological patches in the east-west direction. After that, with the increase of the proportion of resistance caused by human activities, the total area of ecological barrier points gradually decreased. In addition, the maximum recovery value of ecological barrier points showed a downward trend, which may be related to the narrowing of the resistance surface threshold range shown in Fig. 5.

This study identifies critical ecological reserves in four scenarios and illustrates that human activity has a significant impact on the distribution of critical ecological reserves in and around urban centers; and has a minor impact on areas far from urban centers. Therefore, when we construct the ecological pattern,

1. For areas far from urban centers, we can refer to the distribution of critical ecological protection areas when the natural landscape resistance and the resistance of human activities are equal to 1:0 or 2:1.
2. Areas around town centers: We can refer to the distribution of critical ecological protection areas concerning the natural landscape resistance and human activity resistance equal to 1:1.
3. For urban centers: we can refer to the distribution of critical ecological protection areas when the natural landscape resistance and the resistance of human activities are equal to 1:2.

The construction and protection of these critical ecological reserves can connect scattered ecological sources, help reduce the impact and threat of urban construction on local ecosystems and biodiversity conservation, and maintain the integrity and stability of local ecosystems. More importantly, we consider the impact of human activities and natural landscape, which is more conducive to promoting regional ecological protection and the harmonious development of man and nature.

